# Assessing the feasibility of a new approach to measure the full spectrum of CSF dynamics within the human brain using MRI: insights from a simulation study

**DOI:** 10.1101/2024.11.28.625668

**Authors:** E.C. van der Voort, M.C.E. van der Plas, J.J.M. Zwanenburg

## Abstract

Cerebrospinal fluid (CSF) dynamics are essential in waste clearance of the brain. Disruptions in CSF flow are linked to various neurological conditions, highlighting the need for accurate measurement of its dynamics. Current methods typically capture limited aspects of CSF movement or focus on a single anatomical region, presenting challenges for comprehensive analysis. This study proposes a novel approach using Displacement Encoding with Stimulated Echoes (DENSE) MRI to assess the full spectrum of CSF motion within the brain. Through simulations, we evaluated the feasibility of disentangling distinct CSF motion components, including heartbeat- and respiration-driven flows, as well as a net velocity component due to continuous CSF turnover, and tested the performance of our method under incorrect assumptions about the underlying model of CSF motion. Results demonstrate that DENSE MRI can accurately separate these components, and reliably estimated a net velocity, even when periodic physiological motions vary over time. The method proved to be robust for including low frequency components (LFO), incorrect assumption on the nature of the net velocity component and missing CSF components in the model. This approach offers a comprehensive measurement technique for quantifying CSF dynamics, advancing our understanding of the relative role of various drivers of CSF dynamics in brain clearance.

## Introduction

Cerebrospinal fluid (CSF) plays a crucial role in maintaining brain homeostasis by clearing metabolic waste and regulating intracranial pressure.(Hladky & Barrand, 2014; Spector et al., 2015) Disruptions in CSF dynamics have been linked to various neurological conditions, including Alzheimer’s disease, where impaired clearance contributes to the buildup of brain metabolic waste,(Nedergaard & Goldman, 2020; Schubert et al., 2019; Tarasoff-Conway et al., 2015) and hydrocephalus, where abnormal CSF accumulation in the ventricles leads to a range of symptoms.(Reeves et al., 2020; Ringstad et al., 2017) Understanding CSF dynamics is key to unraveling the exact mechanisms underlying these disease processes.

Several physiological factors have been identified to be responsible for the driving of CSF. One such factor is the heartbeat; the inflow of arterial blood induces pulsations of the brain tissue, which generate a pulsatile flow of CSF.(Mestre et al., 2018; Sloots et al., 2021) The Monro-Kellie doctrine states that this inflow of blood requires a compensatory outflow of CSF to preserve a constant intracranial volume and pressure.(Mokri, 2001) Another driver of CSF flow is the respiratory cycle, where changes in venous blood flow similarly induce CSF motion in accordance with the Monro-Kellie principles. In recent years, low frequency oscillations (LFO), with frequencies typically around 0.1 Hz, have also been recognized as a driver responsible for CSF flow, (Wang et al., 2022; Yang et al., 2022) These LFO arise from various sources including spontaneous smooth muscle contractions, also known as vasomotion, (Haddock & Hill, 2005; van Veluw et al., 2020) neuronal activation (e.g., through visual stimulation)(Williams et al., 2023) and fluctuations in arterial CO_2_ levels (Vijayakrishnan Nair et al., 2022), and appear to be more pronounced during non-rapid eye movement (NREM) sleep.(Fultz et al., 2019) Lastly, CSF turnover, due to the continuous excretion of CSF by the choroid plexus, also contributes to the movement of CSF.(Hladky & Barrand, 2014; Smets et al., 2023)

Despite growing interest in CSF dynamics, the relative contributions of these driving factors remain poorly understood and are subject to ongoing debate. Some studies indicate that respiration may be the primary driver of CSF flow,(Dreha-Kulaczewski et al., 2015, 2017; Kollmeier et al., 2022; Yamada et al., 2013) while others emphasize the dominant role of cardiac pulsations.(Daouk et al., 2017; Mestre et al., 2018; Yildiz et al., 2017) Similarly, the extent to which LFO and CSF turnover affect the CSF dynamics is not fully understood, with substantial evidence still lacking. Measuring these CSF dynamics, both qualitatively and quantitatively, is important to understand the role of the various components in CSF dynamics. Several studies have proposed methods to measure CSF dynamics such as by using two-photon microscopy (Iliff et al., 2012; van Veluw et al., 2020) or a range of magnetic resonance imaging (MRI) techniques including arterial spin labelling,(Petitclerc et al., 2021) turbo spin echo,(Hirschler et al., 2019) diffusion weighted imaging,(Wright et al., 2024) phase contrast (PC) MRI,(Matsumae et al., 2014) dynamic contrast enhanced MRI,(Benveniste et al., 2020) blood-oxygenated-level-dependent MRI (Fultz et al., 2019; van der Voort et al., 2024) and balanced steady-state free precession (SSFP)(Wang et al., 2022). However, most of these techniques focus only on one or two specific contributors to the CSF dynamics or limit measurements to a single location, such as the fourth ventricle. Measuring all components simultaneously is challenging due to the difference in time scale and magnitude of these factors. For instance, the net velocity, due to the CSF turnover, is expected to be around 5 μm/s in the subarachnoid space (SAS) based on a crude estimation.^1^ The physiological processes, however, induce periodic motions which are several orders in magnitude larger,(Chen et al., 2015; Takizawa et al., 2017) depending on the location in the brain, complicating concurrent measurements.(Magdoom et al., 2019)

In this study, we propose a new approach to assess the complete spectrum of CSF mobility in a single acquisition, using Displacement Encoding with Stimulated Echoes (DENSE) MRI. This method enables the disentanglement of the different motion components at various locations throughout the brain. We conducted simulations to test the feasibility of this approach with a focus on the reliability of the net velocity as the smallest component. We also assessed and robustness of our method in case of incorrect CSF dynamics model assumptions.

## Methods

### Model of CSF displacement

The CSF dynamics can be described as a linear combinations of distinct motion components. These components include periodic physiological motions, such as the cardiac pulsations, respiration, and LFO’s, alongside a net velocity component. The overall displacement of CSF resulting from these motions can be modeled as follows:

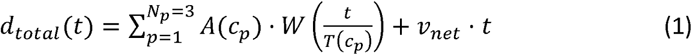

where *A*(*c_p_*) represents the amplitude and 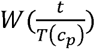 the waveform of the periodic motion, with *T*, being its cycle duration. Both the amplitude, *A,* and cycle duration, *T,* can vary between cycles of the physiological motion, *p* (with *p* = 1,2,3 representing heartbeat, respiration and LFO, respectively, and *c_p_* is the index of the cycles in the physiological process *p*). The time, *t*, is defined as the measurement point, with displacement measured at *t* relative to an arbitrary reference point, which has a CSF displacement of zero by definition. To accurately distinguish between the various motion components, displacement must be measured at a sufficiently large number of independent time points. The unknowns, *A*, *W* and *v_net_*, can be determined using a least squares approach. This method is analogous to the one used in sea-level rise measurements, where the sea-level represents the net velocity component, and lunar tides and wind stress correspond to the periodic physiological processes.(Steffelbauer et al., 2022) Similar to these measurement, the net velocity component of CSF is expected to be much smaller than the periodic motions.

### Measurements of CSF displacement

The CSF displacement can be measured using DENSE, which is an MRI technique that encodes motion in the phase of the MRI signal, somewhat similar to PC MRI.(Adams et al., 2019; Aletras et al., 1999; Sloots et al., 2020) The sequence consists of a motion encoding and motion decoding part, separated by a waiting period called the mixing time (TM) over which displacement is measured. Based on previous work by Sloots et al., we explain the proposed approach using a simplified, single slice DENSE sequence with a single decoding per encoding (Figure 1).(Sloots et al., 2020) This sequence was previously used to measure tissue displacement induced by the heartbeat and respiration.(Sloots et al., 2020, 2021) In order to enable the inclusion of a net velocity component, data should be acquired without cardiac triggering to distinguish between cardiac-induced motion and the net velocity component. The sequence is repeated multiple times in order to cover multiple cycles of the targeted physiological process. These repeats, also known as dynamics, have a fixed repetition time (TR). In the current feasibility study, two different TMs are used which are changed over the dynamics.

**Figure 1.**
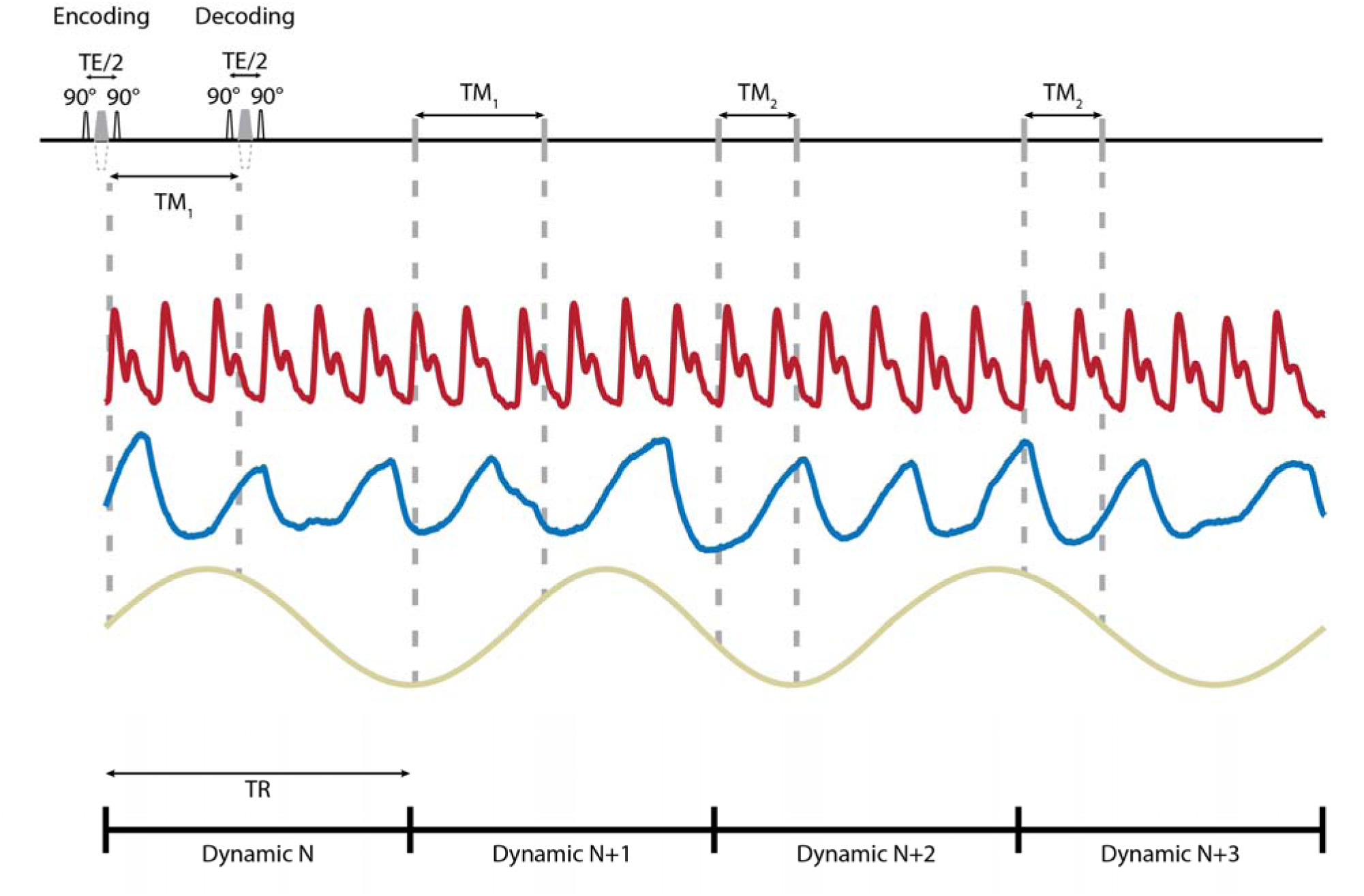
Example of a DENSE measurement and the simultaneously acquired physiological traces. The measurements include a single slice acquisition with a single decoding per TM over four dynamics (repeats) with varying mixing time (TM) over the dynamics. As the DENSE sequence is operated in a non-triggered fashion, the encoding and decoding fall at different positions with respect to the physiological cycles. The motion encoding comprises two 90° radiofrequency pulses with an encoding gradient in between. In the motion decoding part, a single 90° RF pulse is followed by a motion decoding gradient that matches the area of the encoding gradient. TE = echo time, TR = repetition time

The DENSE measurement of CSF displacement that occurs during the TM is particularly sensitive when long TMs are used (Figure 1). Using long TMs is possible as signal loss is predominantly proportional to T1 during the TM and decays with T2 as a function of the echo time (TE). During the TM, displacement of CSF along the direction of the encoding gradient results in phase shift. This phase shift (*φ*) is proportional to displacement according to:

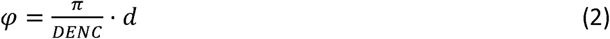

where *d* is the displacement and DENC is the displacement encoding parameter which specifies the displacement required to induce a phase shift of ±π. The lower the DENC value, the higher the motion sensitivity.

Apart from CSF motion, the measured phase includes confounders such as a radiofrequency (RF) offset or a phase induced by off resonance effects, which are independent of the motion sensitizing gradients. In order to separate these confounding factors from motion dependent phase shifts, the polarity of the motion sensitizing gradients is alternated between dynamics. This alternation allows for the isolation of true motion-related phase shifts, and the total measured phase of the MRI signal can then be used to calculate the underlying motion components. As DENSE measures the displacement over the TM, together with a static phase confounder, the measured phase for a single DENSE measurement is given by:

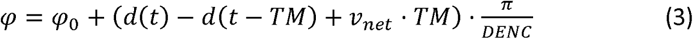

where *φ*_0_ is the static phase offset, independent of motion. Generally, the waveforms of the physiological processes are unknown. Therefore, they are discretized such that they can be estimated from the data. For a single physiological process (*N_p_* = 1), Equation 3 can be approximated as:

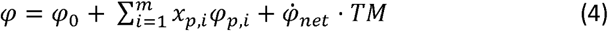

where *φ_p,i_* is the phase accumulation due to the periodic physiological process and 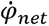 is the phase accumulation due to the net velocity component. The weightings, *x_p,i_*, are essentially binning parameters, reflecting the relative time within the physiological cycle, *t*/*T* at the moment of decoding and encoding. Thus, the physiological motion is retrospectively binned into *m* bins. The relative timings within the periodic physiological cycles can be obtained by simultaneous acquisition of the physiological traces, such as a respiratory belt for the respiration and a pulse oximeter for the cardiac cycle (and, potentially, the LFO’s as well). Similarly, if the relative amplitude of the physiological motion for the cycle at which the measurement are done is known, this can be incorporated in *x_p,i_*, as well. Multiple periodic physiological processes can be added to the equation above as long as weighting *x_p,i_*, is known.

An example of such a recording for the heartrate, respiration and LFO, and the measurements is given in Figure 1. Here, every encoding and decoding fall at different moments with respect to the physiological cycles. For simplification, the corresponding design matrix only for the measurements with respect to the cardiac cycle is shown below. The first column corresponds to the static offset, which is independent of the motion sensitizing gradients. For every measurement, the moment of encoding and decoding are binned into three bins : *φ*_1,1_, *φ*_1,2_ and *φ*_1,3_ (*p* = 1 for the cardiac cycle). The displacement is defined as the position at decoding minus the position at encoding. Therefore the state of encoding gets a minus sign in the design matrix. The sign of the motion sensitizing gradients is alternated between the dynamics, here the second and fourth dynamic have negative encoding gradients.

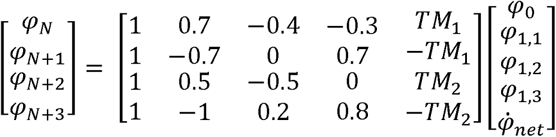

Equation 4 can be solved for the unknowns, *φ*_0_, *φ*_1,*i*_ and 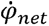, using a least squares approximation if sufficient measurement are available. Because the measurements are non-triggered, the point of encoding has a varying position with respect to the cardiac cycle. Therefore, a reference point was chosen by setting the first bin of the cardiac motion to zero, which essentially removes one column from the design matrix, in this case the second.

### Simulation Design

Simulations were performed for the proposed approach for a single slice DENSE acquisition. Two TMs were used to explore the sensitivity of the data fitting procedure to multiple TMs and the time difference between TMs. The TM was altered between the first and second half of the dynamics. The different motion components were estimated over 60 dynamics with a TR of 6 seconds for every dynamic. The first 15 dynamics were acquired with TM_1_ and positive polarity for the encoding and decoding gradients, the second 15 dynamics with TM_1_ and negative polarity, and the third and fourth 15 dynamics with TM_2_ and alternating gradient polarities, respectively. A total of 60 dynamics were assumed to be sufficient to accurately estimate the different components and would result in a total scan time of 12 minutes, which is acceptable for in vivo scans. The sequence was repeatedly simulated to cover a range of possible combinations of TMs. The TM was varied between 100 and 2500 ms in steps of 100 ms, which resulted in 325 repetitions of the DENSE series.

The displacement for the periodic physiological processes, i.e. the heartbeat, respiration and LFO, were generated from ground truth waveforms. The heartbeat and respiratory waveforms were based on the average of three different in vivo physiological traces (Figure 2). For the LFO, a sinusoidal waveform was used. The amplitude, *A*, and cycle duration, *T*, were varied over time as described by Equation 1 to allow simulating variability in rate and amplitude of the periodic physiological processes. The amplitudes and cycle durations were drawn from normal distributions with mean and SD as follows: durations of the cardiac and respiratory cycle were 1000 ± 100 ms and 5000 ± 1250 ms, respectively, based on the average of available in vivo physiological traces, and the duration of the LFO was set to 10 ± 2 s. The maximum displacement was set to 100 ± 10 µm for the cardiac cycle, 50 ± 15 µm for the respiratory cycle and 75 ± 15 μm for the LFO.

**Figure 2.**
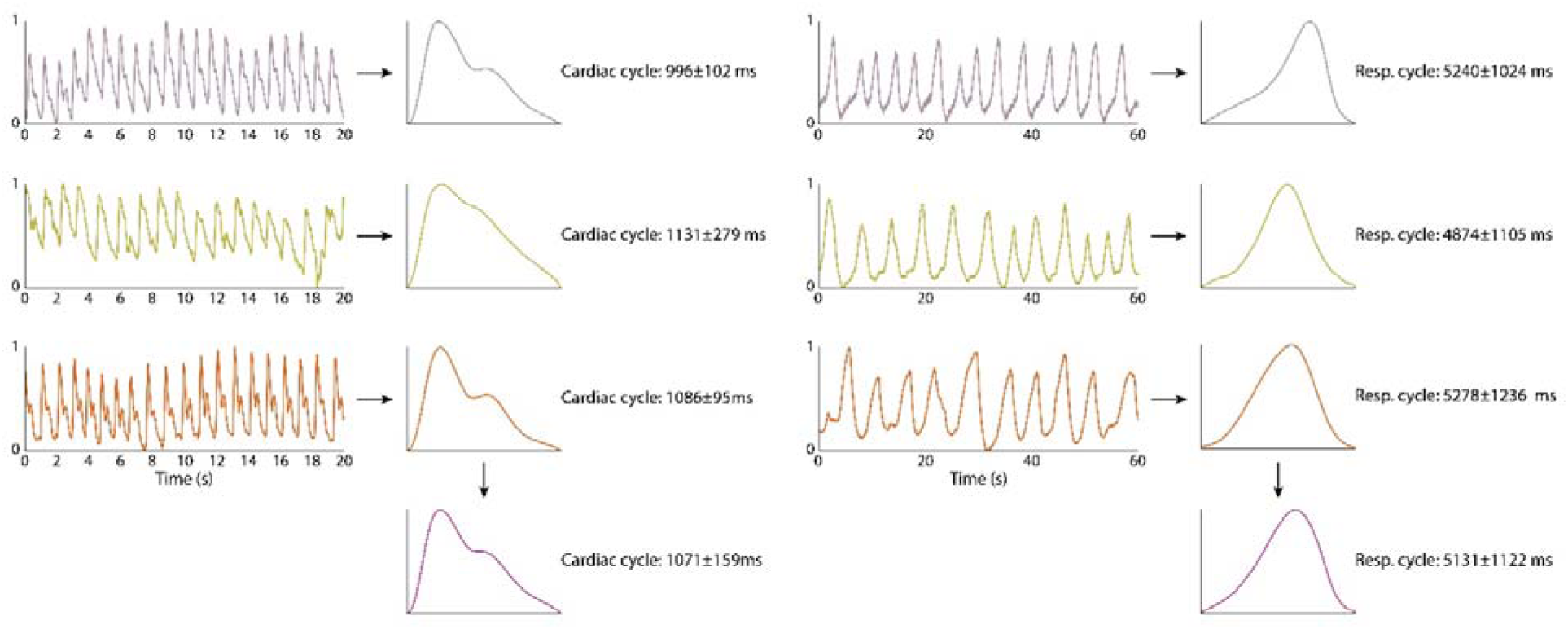
Design of the ground truth for the cardiac (left) and respiratory (right) curves for the simulation data. The ground truth curves were created by first averaging physiological traces of three healthy subjects to obtain representative average cycles of the heartbeat and respiration, respectively. These three averaged cycles are then further averaged to form the ground truth cycles used as basis for the simulations. The duration and amplitude of the cardiac and respiratory cycles are varied for every Monte Carlo run of the simulation using random values from a normal distribution with mean and standard deviation that are derived from those observed in healthy subjects.

The total phase, *φ*, was derived from the ground truth displacement according to Equation 2, using a DENC of 0.125 mm. The measured MRI signal was subsequently generated as complex data *e^iφ^*. Complex noise was added to the signal using a fixed signal to noise ratio (SNR) of 15 for the signal magnitude for all simulations (which is well below realistically achievable SNR with DENSE (Adams et al., 2019)). A Monte Carlo approach was used to evaluate the noise-sensitivity of the estimated parameters with different noise realizations for each Monte Carlo run. A total of 1000 runs was used for each simulated DENSE series (i.e. for each combination of TMs), which is sufficient to estimate the standard deviation (SD) of the results within less than 5% of its true value, with 95% confidence.(Greenwood & Sandomire, 1950) All simulations were implemented in MATLAB 2023b (The MathWorks, Inc., Natrick, MA, USA). The periodic physiological motions were estimated with 10 bins for each process, corresponding to 5%, 15%, 25%, … and 95% of the cycle. As described previously, the first bin, corresponding to the motion at 5% of the cycle, was set the zero. A nonlinear least-squares solver (*lsqnonlin*.m) with default settings was used, with the following start values: 0.1 for the physiological processes, 0.01 for the net velocity component, and 0 for the static offset.

#### Identifying multiple motion components

To assess the feasibility of accurately identifying multiple motion components and estimating the ground truth waveforms when the amplitude and duration are varied over time, simulations were performed for two periodic physiological processes, the heartbeat and respiration, and a net velocity component. The net velocity was set to 5 μm/s.

#### Minimum net velocity

In order to determine the minimal detectable net velocity in the presence of periodic physiological motion, the same simulation was also performed with the net velocity set to 0 μm/s. The 95% confidence interval (CI) of the estimated net velocity was used as a proxy for the sensitivity.

#### Effect of pulsatile net velocity

The arachnoid villi have been referred to as “one-way valves” which open under elevated intracranial pressure. This pressure difference between the arachnoid villi and the superior sagittal sinus will lead to the outflow of CSF into the bloodstream.(Proulx, 2021) This could for instance happen due to the inflow of arterial blood, inducing a net velocity of CSF that is limited to the systolic phase of the cardiac cycle only. A simulation was performed with cardiac- and respiration-induced motion where the heartbeat also induced a pulsatile net velocity. This net velocity was modeled as a constant velocity of 10 μm/s during the systolic phase. The parameter estimation was as described above, assuming a constant net velocity.

#### Effect of additional, non-modeled physiological processes

Lastly, to assess the impact of an incorrect underlying model of CSF displacement is incorrect, a similar simulation was performed, this time adding LFO in addition to the cardiac and respiratory motion. The net velocity was again set to 5 μm/s (constant). The DENSE series was simulated with a total of 90 dynamics to match the number of unknowns relative to the number of measurements. Data were analyzed twice: once using all 90 dynamics for the full set of motion components and once using only the first 60 dynamics for only the cardiac- and respiratory-induced motion and a net velocity component, aligning with previous model assumption.

## Results

### Identifying multiple motion components

The estimated net velocity over all Monte Carlo runs for a ground truth constant velocity of 5 μm/s is presented in Figure 3A and shows that the net velocity was estimated without systematic bias. The mean estimated net velocity ranged between 4.87 and 6.39 μm/s (min and max over all simulated combinations of TM). Longer TMs showed more precise net velocity estimations with lower standard deviations. Using at least one TM above 300 ms gave results with standard deviations between 1.21 and 5.18 μm/s (min and max).

**Figure 3.**
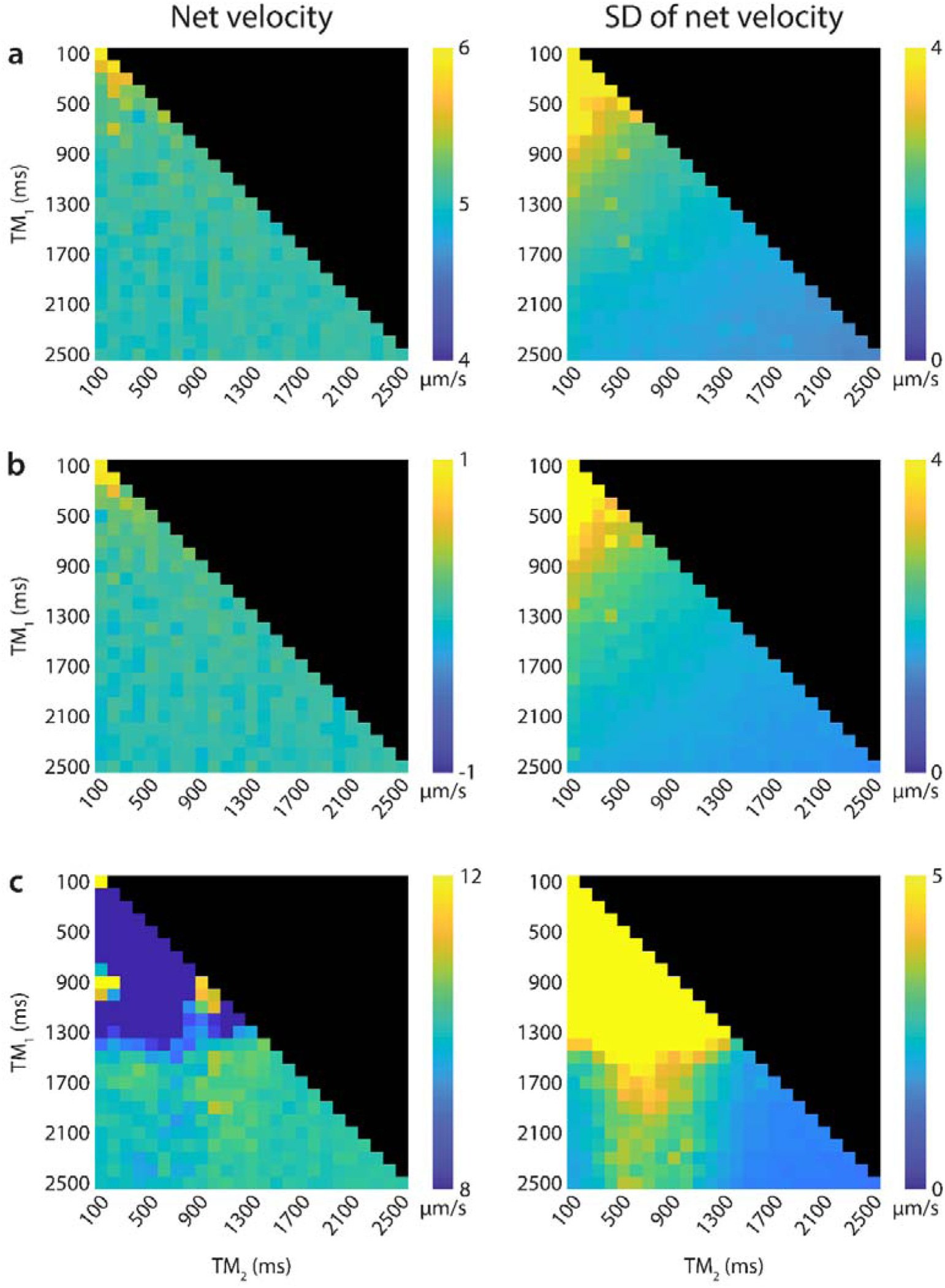
Average estimated net velocity (in µm/s) for various combinations of mixing times (TM) between 100 and 2500 ms, alongside the standard deviation (SD) over all Monte Carlo runs for a simulation including the cardiac and respiratory cycle and a constant ground truth net velocity of 5 μm/s (A) and 0 μm/s (B) and for a pulsatile net velocity of 10 μm/s during the systolic phase of the cardiac cycle (C). The displayed pulsatile net velocity is determined by multiplying the estimated net velocity with the average cardiac period (1000 ms) and dividing it by the average duration of the systolic phase (464 ms). Note that the scales on the color maps vary for optimal visualization of the results, and that the values on the diagonal are obtained with effectively a single TM in the acquisition protocol.

Despite varying amplitude and cycle periods for both heartrate and respiration, the estimated cardiac and respiratory motions, averaged over all TMs and Monte Carlo runs, matched well with the imposed ground truth curves (Figure 4A). As the first bin, which corresponds to 5% of both the cardiac and respiratory cycle, was set to zero to function as a reference point, the overall curves are slightly underestimated compared to the ground truths.

**Figure 4.**
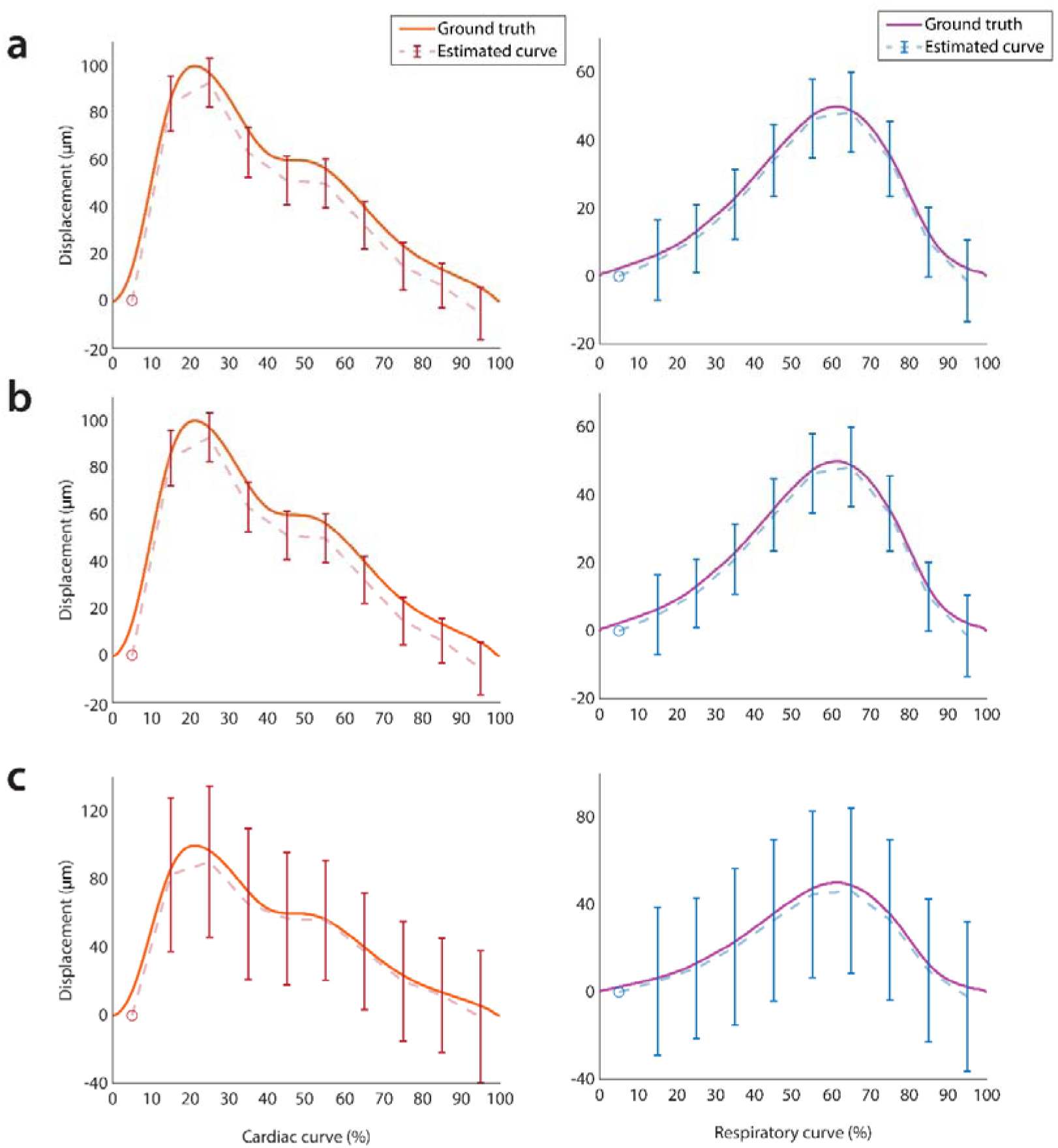
Simulation results showing the estimated cardiac (left) and respiratory (right) waveforms and their ground truths for a simulation with a ground truth constant net velocity of 5 μm/s (A) and 0 μm/s (B) and for a pulsatile net velocity of 10 μm/s during the systolic phase (C). The estimated waveforms were averaged over all TMs and Monte Carlo runs and the error bars indicate the SD over the TMs and Monte Carlo runs. Both waveforms were estimated with 10 bins which correspond to 5%, 15%, 25%, … and 95% of the physiological cycle, where the first bin (5%), indicated by a circle, was used as a semi-arbitrary reference point which has zero displacement an no SD by definition. Both ground truths started at zero at 0% of the cycle, which explains the slight offset between the ground truth and the estimated waveforms.

### Minimum net velocity

Figure 3B shows the net velocity and SD in case where the ground truth velocity was set to 0 μm/s. Similar to previous results, the estimated net velocity is close to the ground truth for most TMs (ranges between −0.12 and 1.12 μm/s, depending on the TM), with simulations performing better when at least one TM was above 300 (max velocity of 0.43 μm/s). The 95% confidence intervals for the cases where effectively one TM was used, i.e. TM_1_ = TM_2_, are shown in Figure 5. The minimal detectable net velocity was approximately 4 μm/s using a TM of 1000 ms and gradually decreased to 2.5 μm/s for a TM of 2500 ms. The estimated physiological curves were similar to the case where the net velocity was 5 μm/s (Figure 4B).

**Figure 5.**
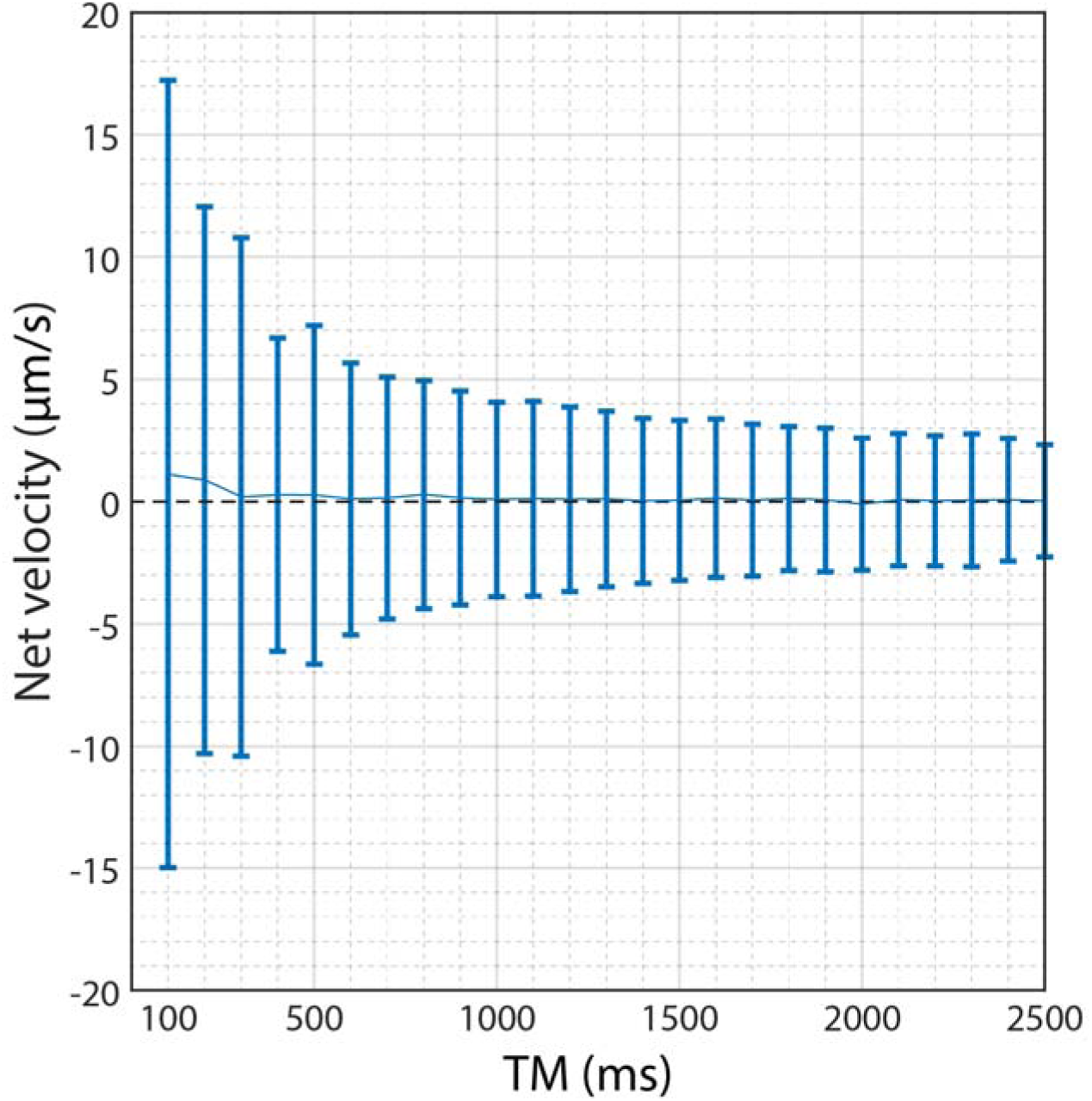
The estimated net velocity (in µm/s), averaged over all Monte Carlo runs, for the cases where TM_1_ = TM_2_, for a simulation including the cardiac and respiratory cycle and a ground truth net velocity of 0 μm/s. The error bars indicate the 95% confidence intervals (1.96 SD). Starting approximately around TM = 1000 ms, the estimated net velocity is close to 0 μm/s. The confidence intervals become progressively smaller for longer TMs. Velocities outside of the 95% confidence interval are regarded as being detectable.

### Effect of pulsatile net velocity

When the ground truth net velocity was pulsatile with the cardiac cycle and only occurred during the systolic phase, the net velocity could only be estimated correctly if at least one TM was approximately equal to or above 1500 ms (covering 1.5 times the average cardiac period), as can be seen in Figure 3C. In this case, the estimated net velocity ranged between 9.00 and 10.90 μm/s (min and max). The estimation of the cardiac- and respiratory-induced motion matched well with the ground truth waveforms but the SDs were larger compared to the simulations with a constant net velocity (Figure 4C).

### Effect of additional non-modeled physiological processes

Adding the LFO as a third physiological process to the model resulted in correct estimation of the net velocity but with considerable dependence on the actual TM, as can be seen in Figure 6A. For TMs below 2100 ms, the estimated net velocity ranged between 3.44 and 5.83 μm/s (min and max). If at least one of the TMs was above 2000 ms, the net velocity was underestimated for most combinations of TM with a mean estimated net velocity of 1.19 μm/s and a range between 0.11 and 5.08 μm/s. The SDs were relatively high, ranging between 2.47 and 11.09 μm/s.

**Figure 6.**
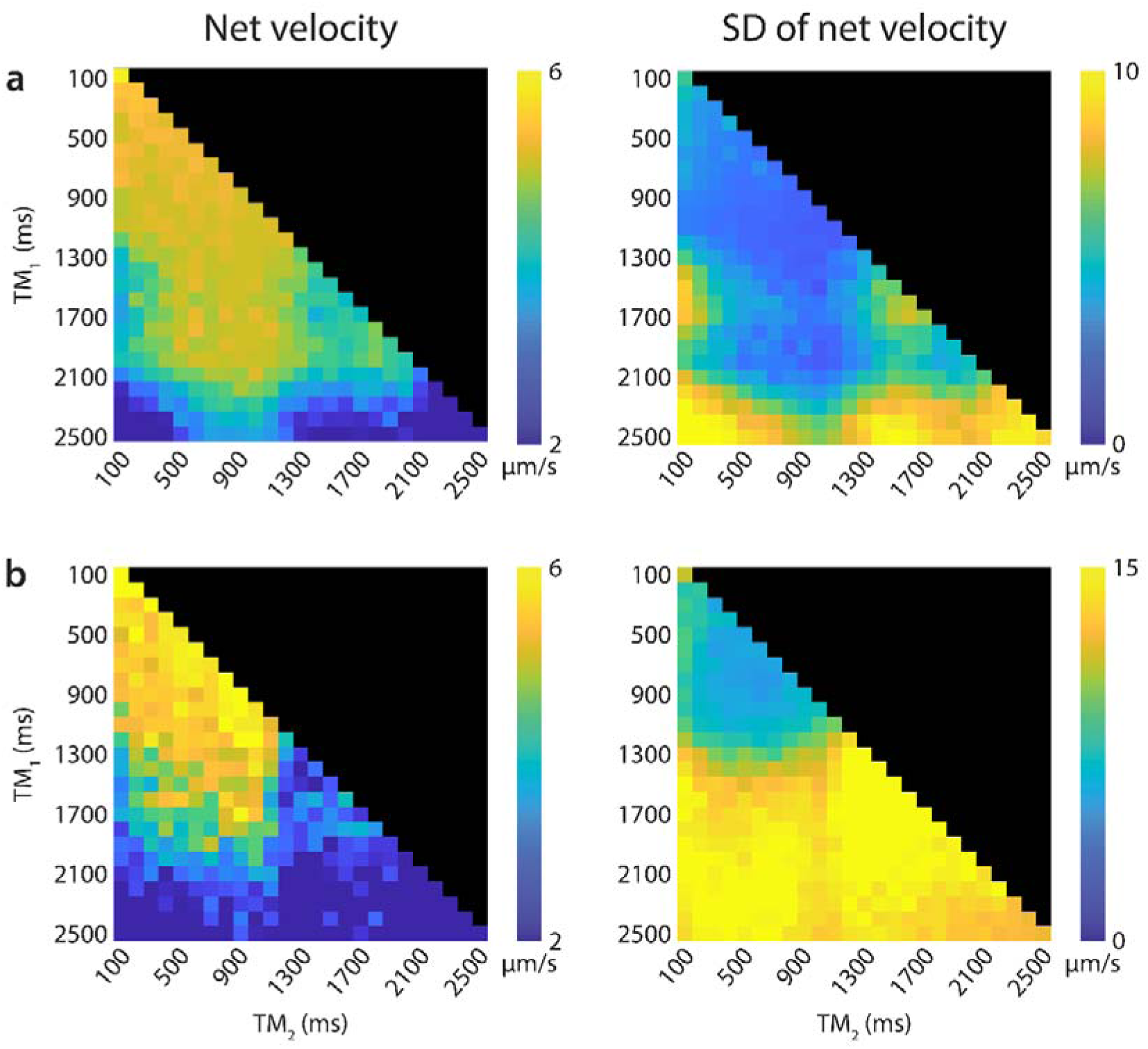
Average estimated net velocity (in µm/s) for various combinations of mixing times (TM) between 100 and 2500 ms, alongside the standard deviation (SD) over all Monte Carlo runs for a simulation including the cardiac-, respiratory- and LFO-induced CSF motion and a ground truth net velocity of 5 μm/s. In case where the LFO was included in the model (A) and be where the LFO was not included (B). Note that the values on the diagonal are obtained with effectively a single TM in the acquisition protocol.

All estimated waveforms of the three physiological processes matched well with the ground truths (Figure 7A). There was an underestimation for all curves which cannot be fully explained by the shift of the first bin. Compared to the simulations without LFO, the SD of the cardiac and respiratory waveforms were larger, as can be seen by the length of the error bars.

**Figure 7.**
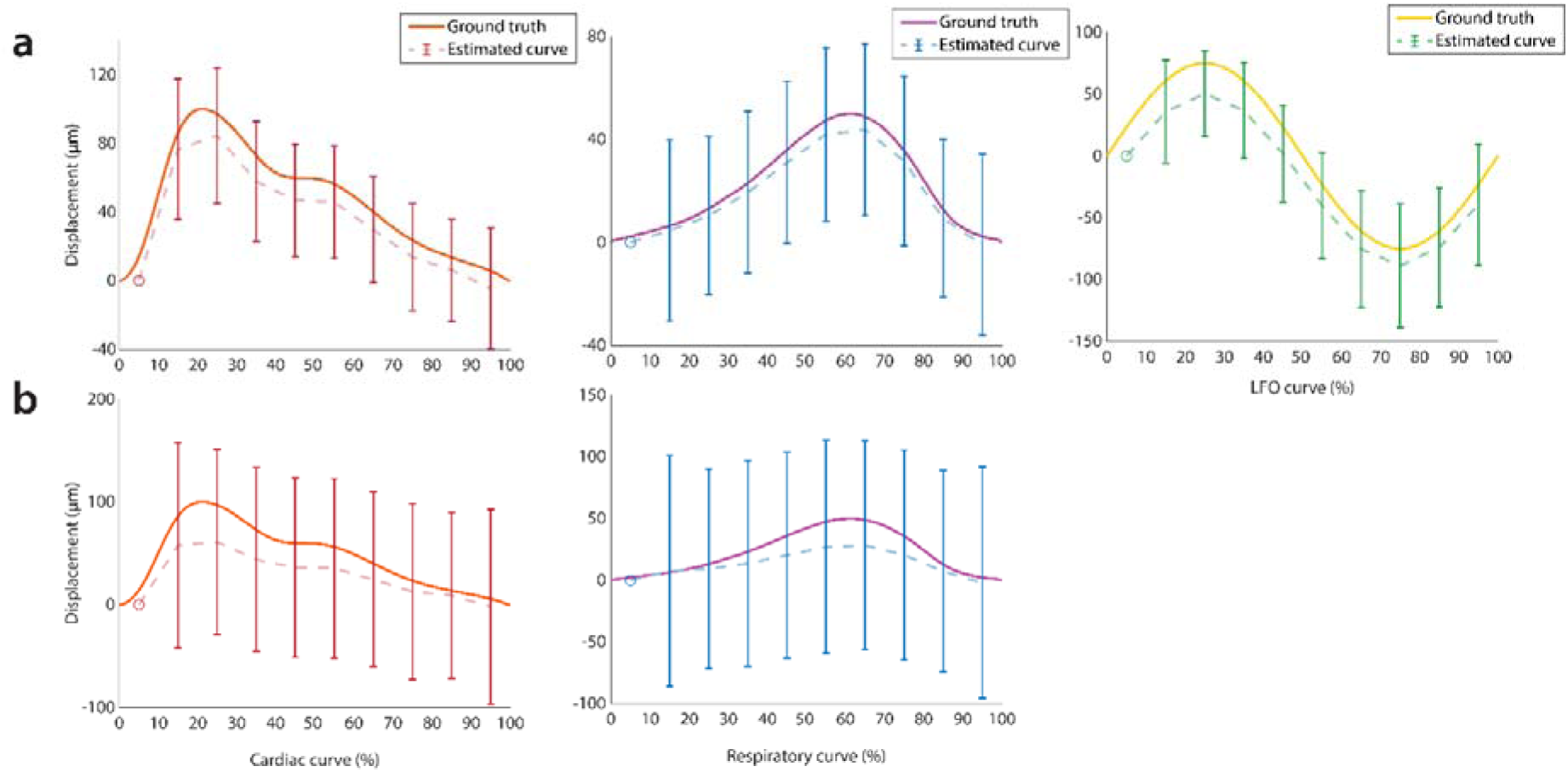
Simulation results showing the estimated cardiac (left), respiratory (middle) and LFO (right) waveforms and their ground truths for a simulation with a ground truth net velocity of 5 μm/s where the LFO was included in the model (A) and excluded (B). The estimated waveforms were averaged over all TMs and Monte Carlo runs and the error bars indicate the SD over the TMs and Monte Carlo runs. All waveforms were estimated with 10 bins which correspond to 5%, 15%, 25%, … and 95% of the ground truth where the first bin (5%), indicated by a circle, was used as a semi-arbitrary reference point which has zero displacement an no SD by definition. Both ground truths started at zero at 0% of the cycle, which partially explains the slight offset between the ground truth and the estimated waveforms.

Incorrect assumption about the underlying CSF model (if the LFO was not taken into account during parameter fitting), did not substantially affect the estimation of the net velocity for shorter TMs (<1500 ms) (Figure 6B). The estimated net velocity showed a similar dependence on the TMs for TMs below 1500 ms compared to the simulation where the LFO was included in the model (Figure 6A). For TMs below 1500 ms, the estimated net velocity ranged between 1.47 and 6.93 μm/s (min and max). When at least one TM was above 1500 ms, the estimated net velocity was on average underestimated (mean of 2.45 μm/s) and ranged between −0.82 and 5.94 μm/s. The cardiac and respiratory curves were underestimated if the LFO was not included in the model, with much larger SDs (Figure 7B).

## Discussion

Measuring the full spectrum of CSF dynamics simultaneously instead of focusing on one or two drivers has the potential to yield better insight into clearance mechanisms. This, however, is difficult because of the wide range in timescales and magnitudes of these drivers. In this study, we tested the feasibility of a new method using DENSE to measure CSF dynamics and a linear least squares approximation to model these dynamics and to disentangle the different contributors. We performed simulations to test the performance of our approach to variability in the CSF dynamics and to incorrect model assumptions. Specifically, we tested how the model performed in cases where contributors were not incorporated or the actual net velocity was in fact pulsatile instead of constant. Our method proved to be capable of accurately disentangling different CSF drivers that have different magnitudes and timescales.

One TM of at least 300 ms or above was needed in order to properly estimate the net velocity, with longer TMs resulting in lower SD. Including, however, LFO as driving factor of the CSF motion, showed a more complex behavior of the accuracy of the estimated net velocity on the TMs. Particularly, for TMs close to 2500 ms (approximately one quarter of the LFO period) appeared to jeopardize correct estimation of the net velocity. This is probably due to the fact that longer TMs enable reasonable amount of displacement due to the net velocity but if these are too long, distinguishing the net velocity from LFO-induced motion becomes complicated. Low frequency components are more difficult to distinguish from a net velocity (which can essentially also be approximated as a very low frequency component) than high frequency components. This will also depend on the net velocity, and the amplitude, waveform and period of the LFO in combination with the distribution of the semi-random measurement points across the LFO-cycle. Current simulations had a maximum TM of 2500 ms, therefore it is unknown what happens when TMs exceed 2500 ms.

If, apart from the net velocity, cardiac and respiratory pulsations were the main physiological motion driving CSF, then, using the current method the minimum detectable net velocity improved with the use of longer TMs. It is important to note that this minimal detectable net velocity – defined as the upper limit of the 95% CI – was estimated over 1000 Monte Carlo runs for a ground truth of zero net velocity. For in vivo estimations, where every voxel can be regarded as an independent measurement, this minimum detectable net velocity scales with 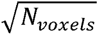 in a region-of-interest based analysis.

In cases where the underlying net velocity was not constant over the measurement duration, but exhibited a pulsatile nature, the net velocity could be correctly estimated if the TM covered at least two systolic phases of the cardiac cycle. The actual estimated velocity was equal to the average of the pulsatile net velocity over the cardiac cycle. This means that our proposed method could yield meaningful net velocity estimates, even though it is ignorant with regard to the true nature of the net velocity (pulsatile or steady).

When a CSF driver, in our case LFO, was not incorporated in the underlying model, but present in the ground truth, the net velocity was still correctly estimated. The estimation of the displacement of the other physiological processes was, however, underestimated, and showed much more variability (larger SDs) which probably depends on TM. This suggests that when the approach is implemented in practice, while no independent trace is available to implement fitting of the LFO component, one should optimize the choice of the TMs by doing further simulations with subject specific characteristics of the heartbeat and respiration frequencies and frequency variability.

All simulations showed that using one TM also gave reliable results. However, in practice, multiple TMs will be useful in order to extend the acquisition to multiple slices and increase the coverage for a given scan duration.

In a previous application of the DENSE sequence to measure brain tissue pulsations,(Sloots et al., 2021) the focus was on cardiac motion. In this study, we used a non-triggered acquisition in order to be equally sensitive to all drivers of CSF. Other studies measuring the CSF dynamics using PC-MRI only focused on one or two physiological contributors (Chen et al., 2015; Takizawa et al., 2017; Töger et al., 2022). Wang et al. used balanced SSFP to measure different timescale drivers simultaneously, but did not take into a net velocity component into account.(Wang et al., 2022) Other methods that did measure the net velocity due to the turnover of CSF, mostly focused on lower regions of brain, i.e. the aqueduct. These studies assume that the contribution of the physiological processes averages out (Huang et al., 2004; Penn et al., 2011; Piechnik et al., 2008). This assumption, however, is questionable for relatively short readouts (Spijkerman et al., 2019). In principle, any partially covered physiological process gives an error in the estimated net velocity, that only decays with O(1/N), where N is the amount of full physiological cycles covered in the measurement duration. In case of respiration-related velocities that are (as in our simulations) an order of magnitude larger than the targeted net velocity, the related error after averaging over 10 respiratory cycles (around 1 minute) is still in the same order of magnitude as the targeted net velocity. Hence, using a maximum-likelihood estimator with a proper model, as proposed here, is much more efficient for obtaining accurate estimations. This also enabled the differentiation of slow net velocities from much higher periodic motions and is similar to the approach used for measuring sea level rise. In that case, the difference in magnitude between the net velocity (sea level rise) and periodic motion (e.g. lunar tides and wind stress) was even larger than for our case, endorsing the applicability of the used method. (Steffelbauer et al., 2022)

### Limitations

Determining the correct magnitude for the different ground truth physiological processes was difficult because the magnitudes differ depending on the brain region. In previous in vivo measurements, we measured a relatively low amplitude of the cardiac and respiratory motion in the peripheral regions of the SAS.(van der Voort et al., 2023) This differs widely from measurements performed in for example the interhemispheric fissure. (Chen et al., 2015) In simulations performed by Wang et al., the heartbeat was modeled as having the highest amplitude, which was six times higher than the respiration and twice as high as the LFO.(Wang et al., 2022) The relative contributions of these physiological processes match well with our simulations. However, their measurements were performed in the fourth ventricle and these relative contributions may also vary between regions.

The heartrate variability is currently implemented by proportionally shortening or lengthening the cardiac waveform over the cardiac period. However, in cases of lower heartrate (i.e. longer intervals), the duration of the diastolic phase is typically more prolonged relative to higher heart rates.(Chung et al., 2004) Additionally, the frequency of the LFO was set at 0.1 Hz, corresponding to vasomotion.(van Veluw et al., 2020) During sleep, however, LFO frequencies as low as 0.05 Hz have been observed.(Fultz et al., 2019) The exact frequency of LFO will affect the dependence on TM for the estimation of the net velocity and these lower frequency components might be more difficult to distinguish from the net velocity. Despite this, using intermediate TMs appears to be effective in accurately estimating the net velocity in presence of LFO, which is advantageous due to the T1 relaxation and motion-related SNR loss for longer TMs for in vivo measurements. Lastly, in vivo aspects such as head motion and magnetic field distortions were not taken into account in the simulations. This is something that will need to be corrected for with e.g. pre-processing of actual in vivo measurements.

## Conclusion

In this study we show the feasibility of a new method, capable of measuring the complete CSF dynamics simultaneously including a net velocity component. This method was shown to be robust for errors in the model assumptions about the underlying CSF dynamics. In order to use this method in vivo, phantom validation is still needed and the optimal choice of the TMs for the acquisition may depend on the frequency of the heartrate, respiration, and the LFOs that are active.

## Acknowledgements

This publication is supported by NWO VICI: Seismology of the brain (18674) and NWO Open Mind: A brainwave inspired by global mean sea-level rise (20696)

## Footnotes

1 For this estimation, the CSF was approximated as a thin shell surrounding a spherical brain with a radius of 65 mm. Taking a CSF excretion rate of 0.30 mL/min and a total CSF volume in this shell of 130 mL,(Huang et al., 2004; Liu et al., 2022) results in a thickness of 2.4 mm and a net velocity of approximately 5 µm/s.

## Notes

### Competing Interest Statement

The authors have declared no competing interest.

## References

Adams, A. L., Kuijf, H. J., Viergever, M. A., Luijten, P. R., & Zwanenburg, J. J. M. (2019). Quantifying cardiac-induced brain tissue expansion using DENSE. Nmr in Biomedicine, 32(2). 10.1002/NBM.4050

Aletras, A. H., Ding, S., Balaban, R. S., & Wen, H. (1999). DENSE: Displacement Encoding with Stimulated Echoes in Cardiac Functional MRI. Journal of Magnetic Resonance, 137(1), 247–252. 10.1006/JMRE.1998.1676

Benveniste, H., Lee, H., Ozturk, B., Chen, X., Koundal, S., Vaska, P., Tannenbaum, A., & Volkow, N. D. (2020). Glymphatic Cerebrospinal fluid and solute transport quantified by MRI and PET imaging. Neuroscience, 474, 63. 10.1016/J.NEUROSCIENCE.2020.11.014

Chen, L., Beckett, A., Verma, A., & Feinberg, D. A. (2015). Dynamics of respiratory and cardiac CSF motion revealed with real-time simultaneous multi-slice EPI velocity phase contrast imaging. NeuroImage, 122, 281–287. 10.1016/J.NEUROIMAGE.2015.07.073

Chung, C. S., Karamanoglu, M., & Kovács, S. J. (2004). Duration of diastole and its phases as a function of heart rate during supine bicycle exercise. American Journal of Physiology. Heart and Circulatory Physiology, 287(5). 10.1152/AJPHEART.00404.2004

Daouk, J., Bouzerar, R., & Baledent, O. (2017). Heart rate and respiration influence on macroscopic blood and CSF flows. Acta Radiologica, 58(8), 977–982. 10.1177/0284185116676655

Dreha-Kulaczewski, S., Joseph, A. A., Merboldt, K. D., Ludwig, H. C., Gärtner, J., & Frahm, J. (2017). Identification of the Upward Movement of Human CSF In Vivo and its Relation to the Brain Venous System. The Journal of Neuroscience⍰: The Official Journal of the Society for Neuroscience, 37(9), 2395–2402. 10.1523/JNEUROSCI.2754-16.2017

Dreha-Kulaczewski, S., Joseph, A. A., Merboldt, K.-D., Ludwig, X.-C., Gärtner, J., & Frahm, X. J. (2015). Inspiration Is the Major Regulator of Human CSF Flow. 10.1523/JNEUROSCI.3246-14.2015

Fultz, N. E., Bonmassar, G., Setsompop, K., Stickgold, R. A., Rosen, B. R., Polimeni, J. R., & Lewis, L. D. (2019). Coupled electrophysiological, hemodynamic, and cerebrospinal fluid oscillations in human sleep. Science (New York, N.Y.), 366(6465), 628. 10.1126/SCIENCE.AAX5440

Greenwood, J. A., & Sandomire, M. M. (1950). Sample Size Required for Estimating the Standard Deviation as a Per Cent of its True Value. Journal of the American Statistical Association, 45(250), 257–260. 10.1080/01621459.1950.10483356

Haddock, R. E., & Hill, C. E. (2005). Rhythmicity in arterial smooth muscle. The Journal of Physiology, 566(3), 645–656. 10.1113/JPHYSIOL.2005.086405

Hirschler, L., Aldea, R., Petitclerc, L., Ronen, I. A., de Koning, P. J., van Buchem, M. A., & van Osch, M. J. (2019). High resolution T2-prepared MRI enables non-invasive assessment of CSF flow in perivascular spaces of the human brain. In Proceedings of the 28th Annual Meeting of ISMRM Montréal, Canada, 2019. Abstract 0746. https://archive.ismrm.org/2019/0746.html

Hladky, S. B., & Barrand, M. A. (2014). Mechanisms of fluid movement into, through and out of the brain: Evaluation of the evidence. Fluids and Barriers of the CNS, 11(1), 1–32. 10.1186/2045-8118-11-26/FIGURES/7

Huang, T. Y., Chung, H. W., Chen, M. Y., Giiang, L. H., Chin, S. C., Lee, C. S., Chen, C. Y., & Liu, Y. J. (2004). Supratentorial cerebrospinal fluid production rate in healthy adults: quantification with two-dimensional cine phase-contrast MR imaging with high temporal and spatial resolution. Radiology, 233(2), 603–608. 10.1148/RADIOL.2332030884

Iliff, J. J., Wang, M., Liao, Y., Plogg, B. A., Peng, W., Gundersen, G. A., Benveniste, H., Vates, G. E., Deane, R., Goldman, S. A., Nagelhus, E. A., & Nedergaard, M. (2012). A paravascular pathway facilitates CSF flow through the brain parenchyma and the clearance of interstitial solutes, including amyloid β. Science Translational Medicine, 4(147). 10.1126/SCITRANSLMED.3003748/SUPPL_FILE/MOVIE_2.MOV

Kollmeier, J. M., Gürbüz-Reiss, L., Sahoo, P., Badura, S., Ellebracht, B., Keck, M., Gärtner, J., Ludwig, H. C., Frahm, J., & Dreha-Kulaczewski, S. (2022). Deep breathing couples CSF and venous flow dynamics. Scientific Reports 2022 12:1, 12(1), 1–13. 10.1038/s41598-022-06361-x

Liu, G., Ladrón-de-Guevara, A., Izhiman, Y., Nedergaard, M., & Du, T. (2022). Measurements of cerebrospinal fluid production: a review of the limitations and advantages of current methodologies. Fluids and Barriers of the CNS 2022 19:1, 19(1), 1–24. 10.1186/S12987-022-00382-4

Magdoom, K. N., Zeinomar, A., Lonser, R. R., Sarntinoranont, M., & Mareci, T. H. (2019). Phase contrast MRI of creeping flows using stimulated echo. Journal of Magnetic Resonance (San Diego, Calif.⍰: 1997), 299, 49–58. 10.1016/J.JMR.2018.12.009

Matsumae, M., Hirayama, A., Atsumi, H., Yatsushiro, S., & Kuroda, K. (2014). Velocity and pressure gradients of cerebrospinal fluid assessed with magnetic resonance imaging: Clinical article. Journal of Neurosurgery, 120(1), 218–227. 10.3171/2013.7.JNS121859

Mestre, H., Tithof, J., Du, T., Song, W., Peng, W., Sweeney, A. M., Olveda, G., Thomas, J. H., Nedergaard, M., & Kelley, D. H. (2018). Flow of cerebrospinal fluid is driven by arterial pulsations and is reduced in hypertension. Nature Communications 2018 9:1, 9(1), 1–9. 10.1038/s41467-018-07318-3

Mokri, B. (2001). The Monro-Kellie hypothesis: applications in CSF volume depletion. Neurology, 56(12), 1746–1748. 10.1212/WNL.56.12.1746

Nedergaard, M., & Goldman, S. A. (2020). Glymphatic failure as a final common pathway to dementia. Science (New York, N.Y.), 370(6512). 10.1126/SCIENCE.ABB8739

Penn, R. D., Basati, S., Sweetman, B., Guo, X., & Linninger, A. (2011). Ventricle wall movements and cerebrospinal fluid flow in hydrocephalus. Journal of Neurosurgery, 115(1), 159–164. 10.3171/2010.12.JNS10926

Petitclerc, L., Hirschler, L., Wells, J. A., Thomas, D. L., van Walderveen, M. A. A., van Buchem, M. A., & van Osch, M. J. P. (2021). Ultra-long-TE arterial spin labeling reveals rapid and brain-wide blood-to-CSF water transport in humans. NeuroImage, 245, 118755. 10.1016/J.NEUROIMAGE.2021.118755

Piechnik, S. K., Summers, P. E., Jezzard, P., & Byrne, J. V. (2008). Magnetic resonance measurement of blood and CSF flow rates with phase contrast--normal values, repeatability and CO2 reactivity. Acta Neurochirurgica. Supplement, 102(102), 263–270. 10.1007/978-3-211-85578-2_50

Proulx, S. T. (2021). Cerebrospinal fluid outflow: a review of the historical and contemporary evidence for arachnoid villi, perineural routes, and dural lymphatics. Cellular and Molecular Life Sciences 2021 78:6, 78(6), 2429–2457. 10.1007/S00018-020-03706-5

Reeves, B. C., Karimy, J. K., Kundishora, A. J., Mestre, H., Cerci, H. M., Matouk, C., Alper, S. L., Lundgaard, I., Nedergaard, M., & Kahle, K. T. (2020). Glymphatic System Impairment in Alzheimer’s Disease and Idiopathic Normal Pressure Hydrocephalus. Trends in Molecular Medicine, 26(3), 285–295. 10.1016/J.MOLMED.2019.11.008/ASSET/B45B85C3-2925-47C9-8413-07252DACF394/MAIN.ASSETS/GR3.JPG

Ringstad, G., Vatnehol, S. A. S., & Eide, P. K. (2017). Glymphatic MRI in idiopathic normal pressure hydrocephalus. Brain, 140(10), 2691–2705. 10.1093/BRAIN/AWX191

Schubert, J. J., Veronese, M., Marchitelli, L., Bodini, B., Tonietto, M., Stankoff, B., Brooks, D. J., Bertoldo, A., Edison, P., & Turkheimer, F. E. (2019). Dynamic 11C-PiB PET Shows Cerebrospinal Fluid Flow Alterations in Alzheimer Disease and Multiple Sclerosis. Journal of Nuclear Medicine⍰: Official Publication, Society of Nuclear Medicine, 60(10), 1452–1460. 10.2967/JNUMED.118.223834

Sloots, J. J., Biessels, G. J., de Luca, A., & Zwanenburg, J. J. M. (2021). Strain Tensor Imaging: Cardiac-induced brain tissue deformation in humans quantified with high-field MRI. NeuroImage, 236. 10.1016/J.NEUROIMAGE.2021.118078

Sloots, J. J., Biessels, G. J., & Zwanenburg, J. J. M. (2020). Cardiac and respiration-induced brain deformations in humans quantified with high-field MRI. NeuroImage, 210, 116581. 10.1016/J.NEUROIMAGE.2020.116581

Smets, N. G., Strijkers, G. J., Vinje, V., & Bakker, E. N. T. P. (2023). Cerebrospinal fluid turnover as a driver of brain clearance. NMR in Biomedicine. 10.1002/NBM.5029

Spector, R., Robert Snodgrass, S., & Johanson, C. E. (2015). A balanced view of the cerebrospinal fluid composition and functions: Focus on adult humans. Experimental Neurology, 273, 57–68. 10.1016/J.EXPNEUROL.2015.07.027

Spijkerman, J. M., Geurts, L. J., Siero, J. C. W., Hendrikse, J., Luijten, P. R., & Zwanenburg, J. J. M. (2019). Phase contrast MRI measurements of net cerebrospinal fluid flow through the cerebral aqueduct are confounded by respiration. Journal of Magnetic Resonance Imaging⍰: JMRI, 49(2), 433–444. 10.1002/JMRI.26181

Steffelbauer, D. B., Riva, R. E. M., Timmermans, J. S., Kwakkel, J. H., & Bakker, M. (2022). Evidence of regional sea-level rise acceleration for the North Sea. Environmental Research Letters, 17(7), 074002. 10.1088/1748-9326/AC753A

Takizawa, K., Matsumae, M., Sunohara, S., Yatsushiro, S., & Kuroda, K. (2017). Characterization of cardiac- and respiratory-driven cerebrospinal fluid motion based on asynchronous phase-contrast magnetic resonance imaging in volunteers. Fluids Barriers CNS, 14, 25. 10.1186/s12987-017-0074-1

Tarasoff-Conway, J. M., Carare, R. O., Osorio, R. S., Glodzik, L., Butler, T., Fieremans, E., Axel, L., Rusinek, H., Nicholson, C., Zlokovic, B. V., Frangione, B., Blennow, K., Ménard, J., Zetterberg, H., Wisniewski, T., & De Leon, M. J. (2015). Clearance systems in the brain-implications for Alzheimer disease. Nature Reviews. Neurology, 11(8), 457–470. 10.1038/NRNEUROL.2015.119

Töger, J., Andersen, M., Haglund, O., Kylkilahti, T. M., Lundgaard, I., & Markenroth Bloch, K. (2022). Real-time imaging of respiratory effects on cerebrospinal fluid flow in small diameter passageways. Magnetic Resonance in Medicine, 88(2), 770. 10.1002/MRM.29248

van der Voort, E. C., Tong, Y., van Grinsven, E. E., Zwanenburg, J. J. M., Philippens, M. E. P., & Bhogal, A. A. (2024). CO2 as an engine for neurofluid flow: Exploring the coupling between vascular reactivity, brain clearance, and changes in tissue properties. NMR in Biomedicine, 37(8). 10.1002/NBM.5126

van der Voort, E. C., van der Plas, M. E. C., & Zwanenburg, J. J. M. (2023). Assessing CSF secretion by measuring net velocity of CSF in the human subarachnoid space using displacement encoding with stimulated echoes at 7T. Proc. Int. Soc. Magn. Reson. Med., 3191.

van Veluw, S. J., Hou, S. S., Calvo-Rodriguez, M., Arbel-Ornath, M., Snyder, A. C., Frosch, M. P., Greenberg, S. M., & Bacskai, B. J. (2020). Vasomotion as a Driving Force for Paravascular Clearance in the Awake Mouse Brain. Neuron, 105(3), 549–561.e5. 10.1016/J.NEURON.2019.10.033

Vijayakrishnan Nair, V., Kish, B. R., Inglis, B., Yang, H. C., Wright, A. M., Wu, Y. C., Zhou, X., Schwichtenberg, A. J., & Tong, Y. (2022). Human CSF movement influenced by vascular low frequency oscillations and respiration. Frontiers in Physiology, 13. 10.3389/FPHYS.2022.940140/FULL

Wang, Y., van Gelderen, P., de Zwart, J. A., Özbay, P. S., Mandelkow, H., Picchioni, D., & Duyn, J. H. (2022). Cerebrovascular activity is a major factor in the cerebrospinal fluid flow dynamics. NeuroImage, 258, 119362. 10.1016/J.NEUROIMAGE.2022.119362

Williams, S. D., Setzer, B., Fultz, N. E., Valdiviezo, Z., Tacugue, N., Diamandis, Z., & Lewis, L. D. (2023). Neural activity induced by sensory stimulation can drive large-scale cerebrospinal fluid flow during wakefulness in humans. PLOS Biology, 21(3), e3002035. 10.1371/JOURNAL.PBIO.3002035

Wright, A. M., Wu, Y. C., Feng, L., & Wen, Q. (2024). Diffusion magnetic resonance imaging of cerebrospinal fluid dynamics: Current techniques and future advancements. NMR in Biomedicine, e5162. 10.1002/NBM.5162

Yamada, S., Miyazaki, M., Yamashita, Y., Ouyang, C., Yui, M., Nakahashi, M., Shimizu, S., Aoki, I., Morohoshi, Y., & McComb, J. G. (2013). Influence of respiration on cerebrospinal fluid movement using magnetic resonance spin labeling. Fluids and Barriers of the CNS, 10(1), 36. 10.1186/2045-8118-10-36

Yang, H. C., Inglis, B., Talavage, T. M., Nair, V. V., Yao, J., Fitzgerald, B., Schwichtenberg, A. J., & Tong, Y. (2022). Coupling between cerebrovascular oscillations and CSF flow fluctuations during wakefulness: An fMRI study. Journal of Cerebral Blood Flow & Metabolism, 42(6), 1091. 10.1177/0271678X221074639

Yildiz, S., Thyagaraj, S., Jin, N., Zhong, X., Heidari Pahlavian, S., Martin, B. A., Loth, F., Oshinski, J., & Sabra, K. G. (2017). Quantifying the influence of respiration and cardiac pulsations on cerebrospinal fluid dynamics using real-time phase-contrast MRI. Journal of Magnetic Resonance Imaging⍰: JMRI, 46(2), 431–439. 10.1002/JMRI.25591

